# Fluctuating DNA methylation sites encode colorectal tumour growth history

**DOI:** 10.64898/2026.06.04.730217

**Authors:** Veselin Manojlovic, Calum Gabbutt, Darryl Shibata, Robert Noble

**Affiliations:** Department of Mathematics, City St George’s, University of London, London, UK; Department of Haematology, Wellcome-MRC Cambridge Stem Cell Institute, University of Cambridge, Cambridge, UK; Centre for Haematology, Department of Immunology and Inflammation, Imperial College London, London, UK; Centre for Evolution and Cancer, Institute of Cancer Research, London, UK; Department of Pathology, University of Southern California Keck School of Medicine, Los Angeles, CA, USA

## Abstract

Determining the nature of human tumour growth is challenging given the impracticality of obtaining detailed data across time. A promising solution is to examine DNA regions whose methylation states fluctuate on clinically relevant timescales, permitting their use as high-resolution lineage tracers. However, existing methods developed for analysing the fluctuating methylation loci of normal tissue and lymphoid cancers are inapplicable to large solid tumours. Here we introduce a mechanistic computational model that tracks the evolution of heritable methylation marks as a tumour grows from a single gland to a mass of many cubic centimetres, and a coupled ABC-SMC inference workflow to estimate tumour growth parameters from multi-region bulk methylation arrays. We applied this framework to data from multiple regions of 10 resected colorectal tumours, including 3 adenomas and 7 carcinomas of diverse sizes and clinical stages. By exploring alternative models, we show that intratumour diversity, in terms of methylation errors, stems more from tumour growth via gland fission than from cell turnover within glands. Moreover, the extent of intratumour diversity varies widely between patients, mainly because of eight-fold variation in gland fission rates but also due to differences in methylation and demethylation rates. Inter-gland divergence patterns are consistent with neutral evolution of colorectal tumours and a cancer stem cell fraction of approximately 1%. As well as helping to resolve the nature of colorectal cancer growth and evolution, our results provide proof of principle for a method that may be adapted to infer the biological parameters of other types of solid tumour.

**Author summary:** Estimating how fast, for how long, and in what way a tumour has been growing inside someone’s body is challenging. We sought a solution by looking at chemical marks on DNA that are imperfectly copied every time a cell divides. We built a computer model to track how the patterns of marks diverge among the cells of a colorectal tumour as it grows. By running many simulations and comparing the outcomes to data from 10 human tumours, we were able to reconstruct how each of them had grown. We found that growth rate varies widely and is not linked to the size of the tumour when it is removed. The rates at which the DNA marks change also vary but to a lesser degree. Our approach provides a new, inexpensive way to reconstruct tumour growth histories from tissue samples collected during routine surgery, without having to take multiple measurements over time. This approach could be applied to other human cancer types.

## Introduction

Human solid tumours are typically observed only once, when resected. Tumour growth parameters, ages, and modes of evolution must therefore be inferred from a single snapshot of a structure that has been growing for months or years. Although clonal lineages can be reconstructed from genetic mutations, somatic DNA methylation errors – heritable marks that accumulate during cell division [1, 2] – potentially offer higher temporal resolution from a widely available, low-cost form of data. Because methylation patterns are copied with small but measurable error rates at each replication, they act as endogenous molecular clocks. Early studies exploited this property to infer tumour ancestry and growth dynamics in colorectal tumours using small panels of CpG (cytosine then guanine) tags [3–5].

More recently, Gabbutt *et al*. identified a class of fluctuating CpG (fCpG) loci whose methylation state is neither constitutively methylated nor unmethylated, but instead fluctuates on clinically relevant timescales due to ongoing methylation and demethylation errors [6]. These fCpG loci serve as high-resolution lineage tracers whose divergence encodes the recent cell division history of the sampled tissue [6, 7]. The subsequent development of the EVOFLUx method demonstrated that fCpG-based inference of evolutionary dynamics – including tumour growth rate, age, and epimutation rates – is feasible at clinical scale from bulk methylation profiles in lymphoid cancers [8].

Colorectal cancer – the third most common cancer globally and a leading cause of cancer-related mortality [9] – provides an ideal system for adapting fCpG inference methods to solid tumours. Colorectal tumours grow by gland (crypt) fission, the neoplastic counterpart of normal crypt renewal [10]. Each tumour gland is maintained by a small population of cancer stem cells (CSCs) with indefinite proliferative capacity [11, 12], analogous to the stem cell compartment in normal colonic crypts [13]. Colorectal tumour glands therefore retain a hierarchical organisation in which a few stem cells generate the bulk of the gland population, and tumour growth at the tissue level is driven by fission of these gland units. Analyses of both genetic and epigenetic diversity suggest that the majority of intra-tumour genetic heterogeneity arises early in the final clonal expansion, and that subsequent inter-glandular evolution is effectively neutral [5, 14, 15]. According to this “Big bang” model, tumour growth history can be inferred from single-time-point spatial data, such as from glands sampled from distant locations in a resected tumour. A growing body of computational biology research uses agent-based models and likelihood-free inference in solid tumour evolution in general [16–20], and colorectal cancer in particular [3, 21–24]. There are however no existing methods for exploiting the rich information contained in spatially-structured fCpG data.

Here we present *methdemon*, an agent-based model of colorectal tumour growth and fCpG methylation array dynamics at the gland level, coupled with *methabc*, an ABC-SMC inference workflow that estimates tumour growth parameters, principally gland fission rates and epimutation rates, from multi-region bulk methylation arrays. We apply the framework to infer the biological parameters of 10 colorectal tumours.

## Results

### Patient cohort and fluctuating CpG methylation data

Methylation arrays from samples of 10 tumours (7 carcinomas, 3 adenomas, spanning a range of sizes and clinical stages; Table 1) were profiled using the Illumina Infinium Methylation EPIC BeadChip platform, yielding beta-value measurements at approximately 850,000 CpG sites per gland sample at very high purity (Methods). Eight glands were sampled from two spatially separated regions of each resected tumour (sides A and B), providing a dataset of 80 gland-level bulk methylation profiles with known spatial structure (Fig 1).

**Table 1.** Patient and tumour characteristics.

| Tumour | Sex | Age | Size (cm) | Stage | Glands (A/B) | Diagnosis |
| --- | --- | --- | --- | --- | --- | --- |
| D | F | 49 | 2.0 | 1 | 2/6 | Carcinoma |
| E | M | 55 | 6.1 | 1 | 4/4 | Carcinoma |
| F | F | 71 | 1.8 | 1 | 4/4 | Carcinoma |
| I | M | 78 | 3.6 | 3 | 4/4 | Carcinoma |
| J | F | 93 | 5.0 | 3 | 4/4 | Carcinoma |
| M | F | 76 | 3.0 | 2 | 4/4 | Carcinoma |
| U | F | 77 | 3.9 | 2 | 4/4 | Carcinoma |
| P | M | 71 | 3.5 | — | 3/5 | Adenoma |
| S | M | 66 | 6.0 | — | 4/4 | Adenoma |
| X | F | 69 | 2.5 | — | 4/4 | Adenoma |

**Fig 1.**
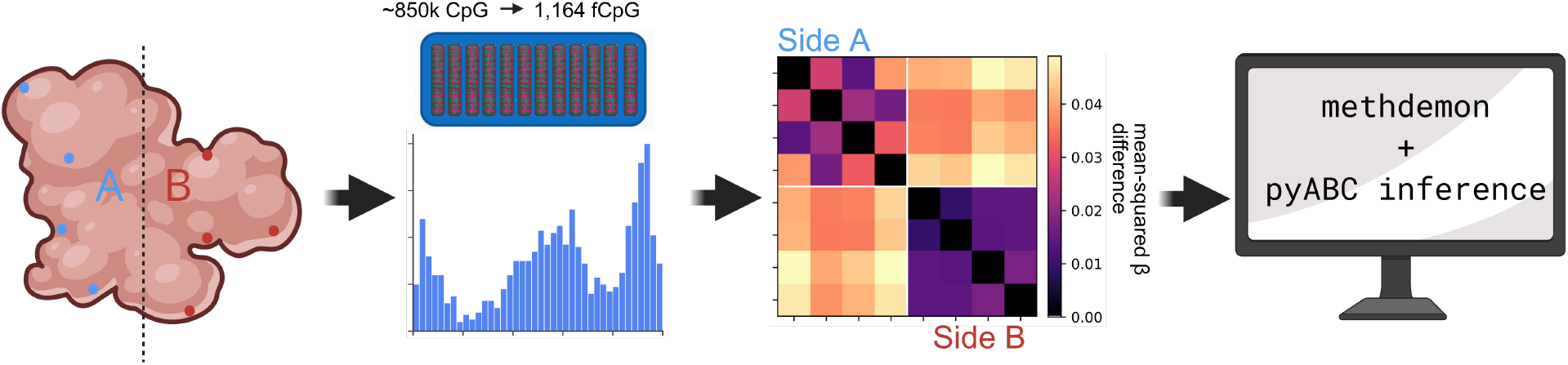
The complete pipeline from sampling to inference. Eights glands worth of bulk methylation arrays are sequenced per tumour, then filtered for fCpG sites and each gland’s fCpG distribution is determined. Their distance matrices are then calculated before they enter the *methabc* inference workflow.

Stochastic gains and losses of methylation at individual CpG sites during somatic cell division produce heritable lineage-informative variation in DNA methylation state [6]. At CpG sites that are neither constitutively methylated nor unmethylated across the population, the methylation fraction of each gland (the *fCpG array*) acts as a cell-intrinsic lineage barcode: genetically identical cells drift apart in methylation state through repeated rounds of division, and the resulting array diversity encodes the number of cell divisions since the most recent common ancestor. These *fluctuating CpG* (fCpG) loci have been used to reconstruct lineage histories in normal tissues and in cancer [6–8].

To identify fCpG loci we followed a procedure similar to that of Gabbutt *et al*. [8], using methylation data from 138 colorectal tumour samples in The Cancer Genome Atlas (TCGA) as a reference cohort (Methods). This procedure identified 1,165 fCpG loci whose methylation states carry information about the history of cell divisions rather than cell type identity. Across the cohort, per-gland fCpG arrays showed the characteristic W-shaped distribution of site beta values (Fig 2A, 2B), consistent with epimutation rates in the intermediate regime where ancestral methylation states are partially but not fully overwritten.

**Fig 2.**
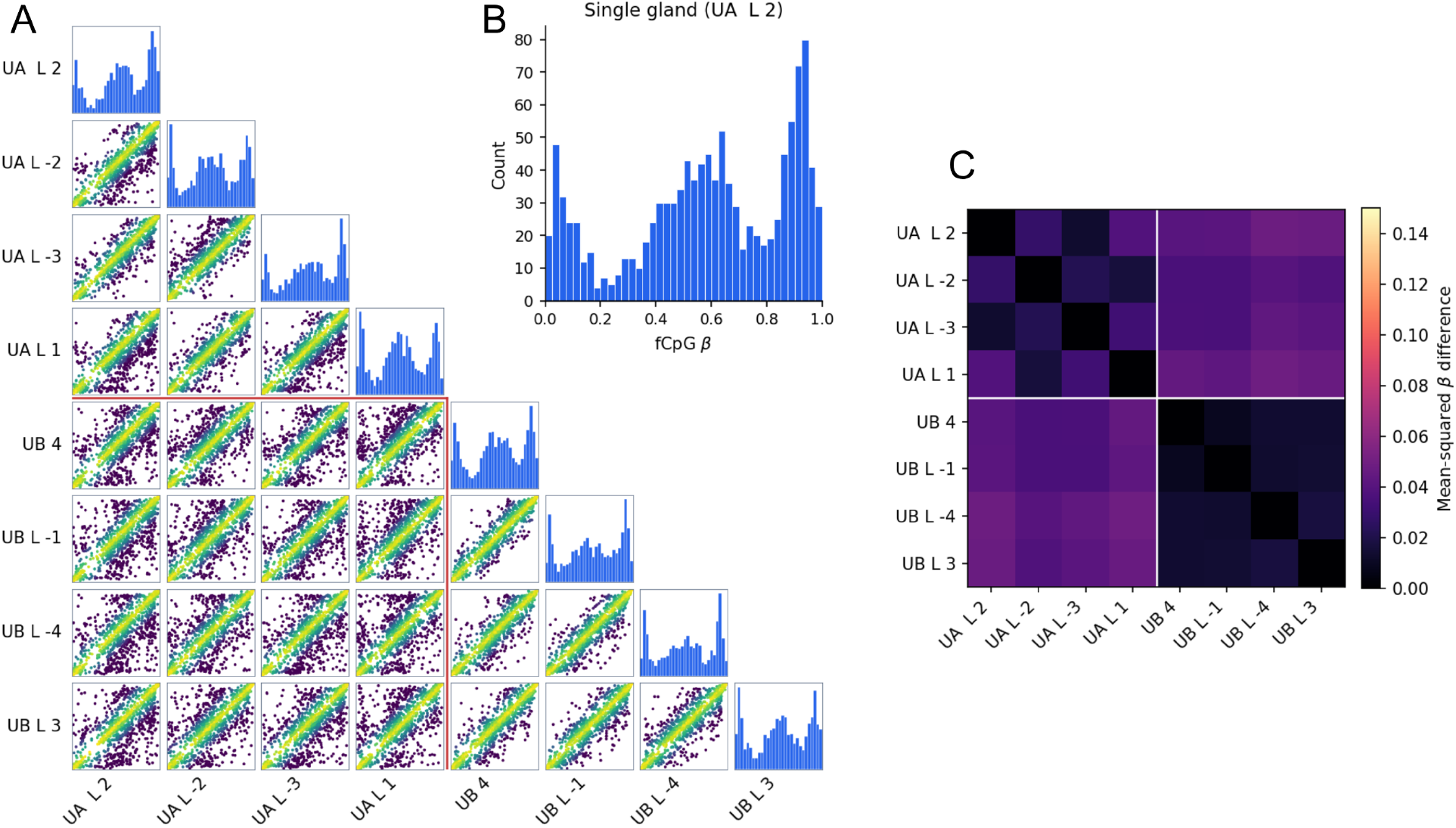
Multi-region bulk fCpG methylation data (tumour U, *n* = 8 glands, 4 per side). **A:** Lower-triangular matrix of paired gland *β* densities; diagonal shows each gland’s univariate *β* histogram, lower triangle shows the 2D density of paired *β* values between glands (darker = higher density). Red lines mark the boundary between side A (top left 4 glands) and side B (bottom right 4 glands). **B:** Single-gland fCpG *β* distribution showing the characteristic W-shape, with peaks near 0, 0.5, and 1. **C:** Inter-gland distance matrix (mean squared difference in *β* across fCpG loci); the 4-by-4 block structure separating the two sides is clearly visible.

### Spatial separation predicts fCpG array divergence in multi-region colorectal cancer samples

To quantify inter-gland methylation divergence, we computed the inter-gland distance matrix **D** for each tumour, defined as the mean squared difference in fCpG methylation fraction between all pairs of sampled glands (Eq 3; Methods). Under the hypothesis that glands from the same side of the tumour diverged from a common ancestor more recently than glands from opposite sides, within-side distances should be systematically lower than between-side distances.

This prediction was confirmed across the full cohort. The inter-gland distance matrix for each tumour showed a clear block structure: gland pairs from the same side consistently are more similar than pairs from opposite sides (Fig 2C, S1). Across all 10 tumours, the median within-side pairwise distance was significantly lower than the median between-side distance (Fig 3). Every tumour exhibited the expected pattern, with between-to-within distance ratios ranging from 1.12 (tumour D) to 4.71 (tumour I). Tumours with higher ratios (I, U, M) tended to show clearer hierarchical block structure in their distance matrices, consistent with a larger gap between within-side coalescence times and the coalescence time of the whole tumour. The weaker separation observed in tumour D (ratio 1.12) may reflect its unusual sampling configuration (2 glands from side A, 6 from side B), which prohibits precise estimation of within-side distance on the minority side.

**Fig 3.**
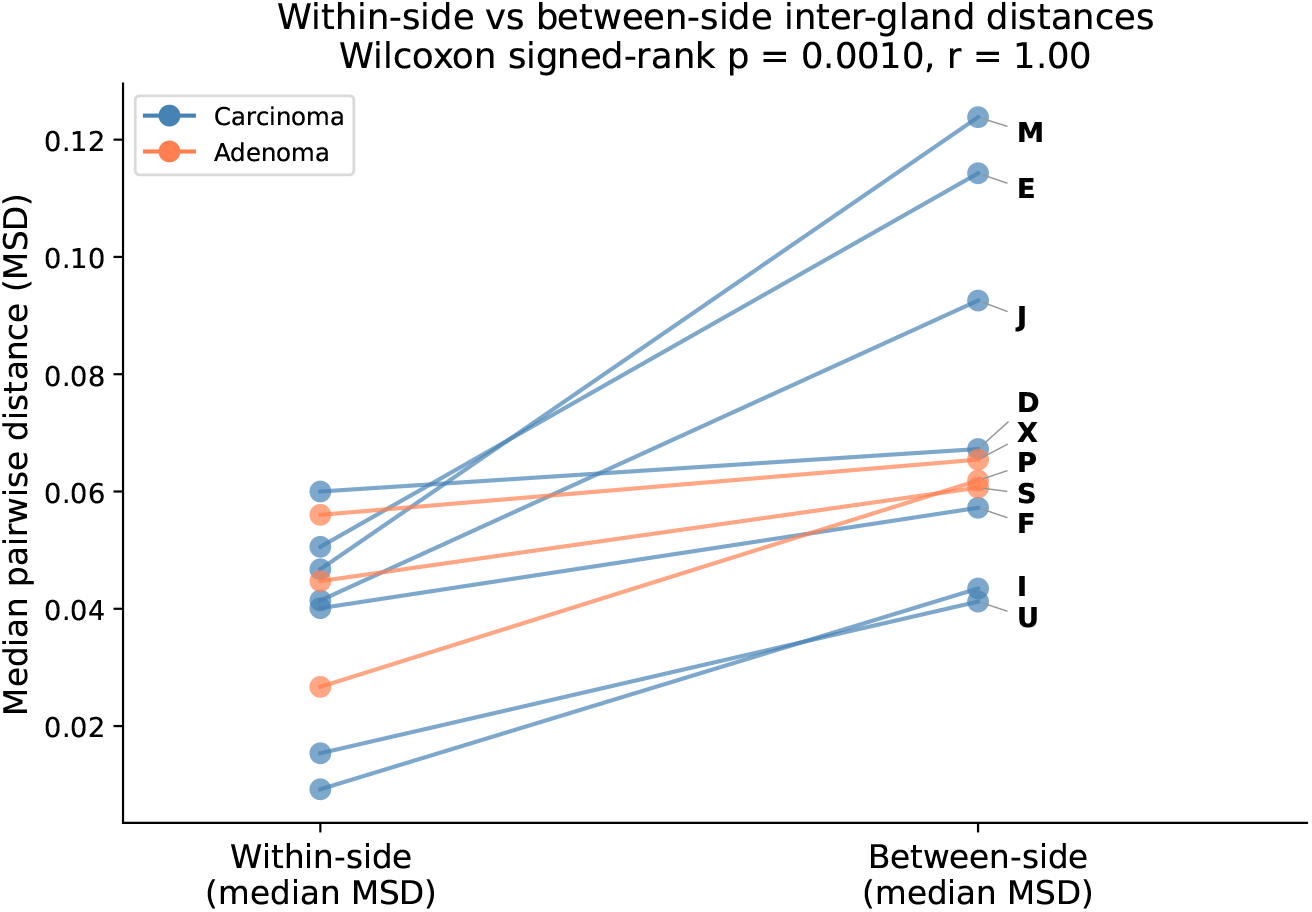
Within-side versus between-side inter-gland distances across the cohort. For each tumour, the median within-side pairwise distance is plotted against the median between-side distance. All 10 tumours have a higher between-side distance than within-side.

The magnitude of inter-gland distances varied substantially across patients (Fig S1), suggesting heterogeneity in either the growth history or the epimutation rates of individual tumours. We reasoned that this variation encodes information about tumour-specific evolutionary parameters recoverable by a calibrated inference framework, motivating the development of the model described in the following section.

### A calibrated agent-based model recapitulates multi-gland fCpG dynamics

To infer tumour-specific growth parameters from the observed fCpG data, we developed *methdemon*, an agent-based model of colorectal tumour growth and fCpG methylation dynamics. In this model, each gland is represented as a deme: a local, well-mixed subpopulation that grows and reproduces as a unit. The model simulates tumour expansion as a branching process of gland fissions, with each gland modelled as a deme of *K* = 100 cancer stem cells undergoing stochastic methylation and demethylation at fCpG loci (Fig 4A). Before performing inference, we characterised how each model parameter shapes the summary statistics to determine which are identifiable from fCpG array data. We ran *methdemon* simulations for a Latin hypercube sample of 200 parameter sets drawn from our inference prior (Table 2), with all other settings matched to the inference configuration (8 sampled glands, 2 sides, *K* = 100, *F* matched to cohort-average tumour size).

**Table 2.** Prior distributions for ABC inference.

| Parameter | Prior ($\log_{10}$ scale) | Original scale range |
| --- | --- | --- |
| $\log_{10} \gamma_{\text{meth}}$ | $\mathcal{U}(-4, -2)$ | $[10^{-4}, 10^{-2}]$ |
| $\log_{10} \gamma_{\text{demeth}}$ | $\mathcal{U}(-4, -2)$ | $[10^{-4}, 10^{-2}]$ |
| $\log_{10} \phi$ | $\mathcal{U}(-3.3, -1)$ | $[5 \times 10^{-4}, 10^{-1}]$ |
| $\log_{10} \mu$ | $\mathcal{U}(-5, -2)$ | $[10^{-5}, 10^{-2}]$ |
| $s$ | $\mathcal{U}(0, 0.2)$ | $[0, 0.2]$ |

All rate parameters were sampled on a $\log_{10}$ -transformed scale. $\mathcal{U}(a, b)$ denotes a uniform distribution on $[a, b]$ .

**Fig 4.**
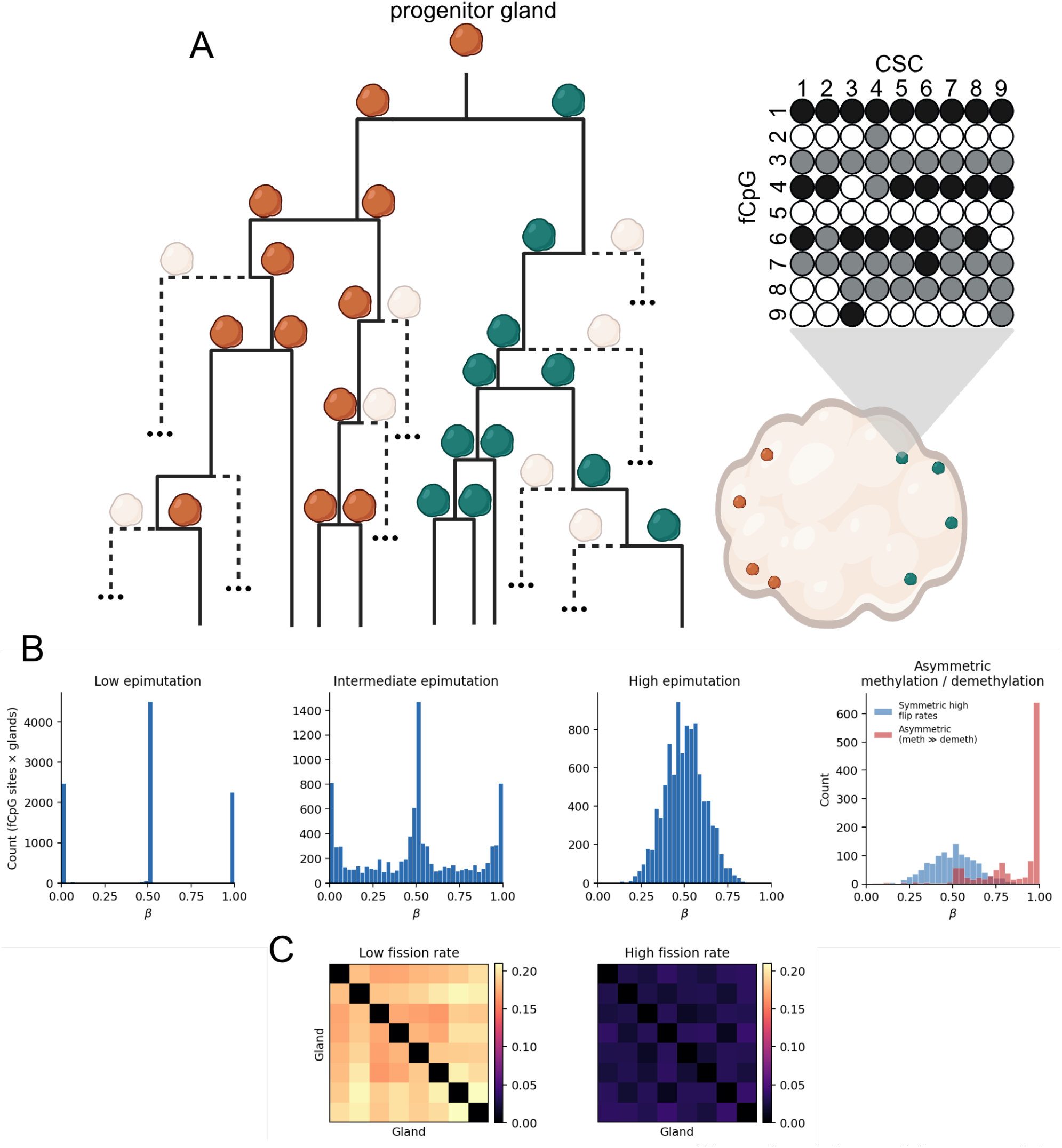
Agent-based model structure and example output. **A:** Hierarchy of the *methdemon* model; only the glands which are sequenced in the end are tracked in the simulations, with the rest discarded during the run. Each gland has a well-mixed population of CSCs inside it, and each CSC has its own distinct fCpG array. **B:** The influence of epimutation rates on individual fCpG arrays; per-gland fCpG *β* distributions (pooled across 8 glands) under four epimutation regimes. Low epimutation rates produce bimodal peaks at 0 and 1; intermediate rates yield the characteristic W-shape; high rates collapse to a unimodal peak whose location depends on the methylation/demethylation ratio. **C:** the inter-gland squared-difference distance matrices at low and high fission rates, illustrating how fission controls the number of cell divisions separating glands and thus the magnitude of inter-gland divergence.

The parameter sweep revealed how epimutation rates *γ*_meth_ and *γ*_demeth_ (per cell division) jointly control the shape and bias of the per-gland fCpG distribution (Fig 4B). At low combined epimutation rates, fCpG sites retain their founder state and the distribution is sharply bimodal, such that all *β* values are close to 0 or 1. At high rates, repeated overwriting of the founder state drives the distribution towards a unimodal peak near the ratio *γ*_meth_*/*(*γ*_meth_ + *γ*_demeth_). At intermediate rates the characteristic W-shape emerges, in which most sites remain near 0 or 1 but a subset has drifted to intermediate values.

The fission rate *φ* controls the per-gland rate at which new glands are produced during expansion and thus the number of cell divisions separating sampled glands. Higher *φ* compresses tumour expansion into fewer cell divisions and reduces inter-gland fCpG divergence, directly shaping the magnitude and block structure of the inter-gland distance matrix (Fig 4C). Because the per-gland fission rate enters the dynamics only through the product *φK*, the fission rate and carrying capacity are structurally non-identifiable: rescaling *K* by a factor *c* is exactly compensated by rescaling *φ* by 1*/c*. We confirmed this degeneracy empirically with a sensitivity analysis. Distance matrices at *K* = 100 and *K* = 1, 000 differed by a Frobenius norm comparable in magnitude to varying other parameters within the prior. A finite-size regime emerges at *K* = 10, where deme capacity approaches the number of founding lineages and additional drift appears. We therefore fixed *K* = 100 for inference and interpret *φK*, rather than *φ* alone, as the biologically meaningful quantity.

Sensitivity analyses established that, while epimutation and fission rates are identifiable from the inter-gland distance matrix (Methods; Fig S2), the driver mutation rate *µ* and selective advantage *s* leave only weak, mutually confounded signatures. Because driver mutations confer a multiplicative fission advantage, increasing either *µ* or *s* produces similar, modest perturbations that are dominated by the stronger signals of fission and epimutation. Consequently, although these parameters were retained in the model for structural completeness, their marginal posteriors closely track their priors and are not reported as biological findings.

### Colorectal tumour glands grow as a near-pure branching process

We applied the ABC-SMC inference workflow to each tumour in the cohort independently to obtain posterior distributions for the five model parameters (Methods; Table 2). The inference framework succeeded in reproducing key qualitative features (compare Fig 5 to Fig 2) and quantitative aspects of the data (Fig S3; Methods). These inference results suggest that the data are broadly consistent with the assumption underlying our agent-based model, that colorectal tumours grow through successive gland fissions from a single progenitor. Real tumours, however, may undergo a period of post-expansion steady-state turnover in which cell birth and death continue within glands at carrying capacity without further fission. A key question is whether such a turnover phase would substantially alter the inter-gland fCpG divergence patterns on which inference relies.

**Fig 5.**
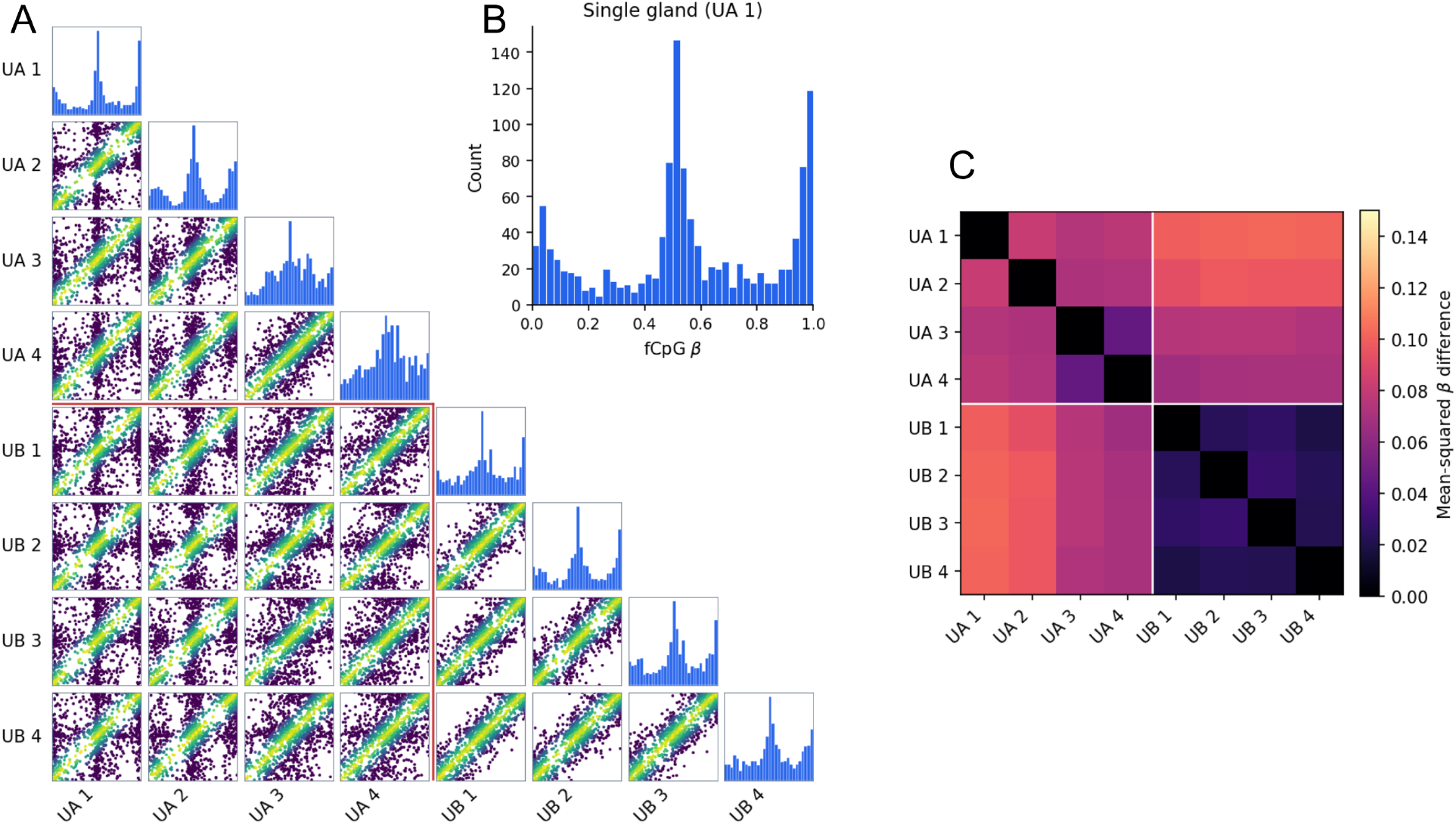
Simulated multi-region fCpG arrays based on the outputs of a single run of *methdemon*. **A:** Paired-gland *β* densities. **B:** Single-gland fCpG *β* distribution. **C:** Inter-gland distance matrix. Parameters were drawn from the posterior distributions of tumour U, as shown in Fig 2.

Two features of the data indicate that the expansion phase dominates the inter-gland divergence signal. First, the clear block structure in the inter-gland distance matrices (Fig 2C; Fig S1) reflects a hierarchical branching history: glands from the same side of the tumour are consistently more similar than glands from opposite sides, as expected if fCpG divergence accumulated primarily during the branching process that produced the spatial separation. Because steady-state turnover acts symmetrically on all glands regardless of their spatial relationships, it would tend to erode the hierarchical structure emerging due to branching. The pronounced block structure across the cohort therefore suggests that expansion is the dominant contributor to the observed signal.

Second, the fCpG distributions within individual glands retain a characteristic W-shape (Fig 2A, 2B), indicating that epimutation rates lie in the intermediate regime where ancestral methylation states are partially but not fully overwritten. Extended turnover at carrying capacity would progressively drive within-gland distributions towards a unimodal distribution centred at 0.5 through continued neutral drift and epimutation.

To quantify the effects of post-expansion turnover, we performed a sensitivity analysis comparing simulated distance matrices across turnover fractions (0%, 15%, 30%, 50%) over a grid of 200 parameter sets sampled from the prior (Methods). Turnover modestly increased overall divergence (median mean pairwise distance +30% at 50% turnover; Fig S4A) but progressively harmonised the between-side and within-side block structure (median ratio: 1.14 at 0% to 1.04 at 30%; Fig S4B, S4C). The observed between/within ratios in the cohort (median 2.24) well exceed the median in the simulations even for 0% turnover, suggesting that expansion is the main driver of the array divergence.

### Inferred gland fission rates vary widely across patients

Applying the ABC-SMC inference framework to all 10 tumours yielded posterior distributions for each tumour’s fission rate, methylation rate, and demethylation rate (Table 3). The posterior distributions of the remaining model parameters – that is, driver mutation rate and selective advantage – stayed broad across all tumours, consistent with effectively neutral inter-glandular dynamics during expansion, and these parameters are therefore excluded from further interpretation.

**Table 3.** Posterior summary statistics for all tumours.

| Tumour | Fission rate $\phi$<br>($\times 10^{-3}$) | Methylation rate $\gamma_m$<br>($\times 10^{-3}$) | Demethylation rate $\gamma_d$<br>($\times 10^{-3}$) |
| --- | --- | --- | --- |
|  | median [90% CI] | median [90% CI] | median [90% CI] |
| D | 4.8 [1.5, 31] | 1.9 [1.0, 3.2] | 2.0 [1.0, 3.3] |
| E | 2.8 [0.97, 16] | 2.3 [1.3, 4.0] | 2.2 [1.2, 3.7] |
| F | 8.9 [2.2, 48] | 3.0 [1.7, 5.0] | 1.6 [0.91, 2.8] |
| I | 14 [2.7, 95] | 4.8 [2.3, 7.9] | 0.79 [0.18, 2.0] |
| J | 3.6 [0.90, 29] | 1.2 [0.57, 2.3] | 1.8 [0.98, 3.8] |
| M | 2.2 [0.67, 17] | 1.1 [0.64, 2.0] | 2.0 [1.2, 3.3] |
| U | 17 [1.6, 82] | 2.8 [1.5, 4.5] | 1.7 [0.89, 3.0] |
| P | 11 [2.5, 44] | 0.98 [0.49, 1.9] | 3.0 [1.6, 5.3] |
| S | 6.8 [1.6, 37] | 1.0 [0.48, 2.3] | 3.3 [1.6, 6.4] |
| X | 4.5 [1.3, 23] | 0.75 [0.27, 1.9] | 2.6 [1.3, 4.9] |

Median and 90% credible intervals from the final-generation ABC-SMC posterior. All rates are on the natural scale (cell<sup>-1</sup> cell div<sup>-1</sup> for fission rate; cell div<sup>-1</sup> site<sup>-1</sup> for epimutation rates).

Median inferred per-cell fission rates spanned a factor of approximately eight, or almost a full order of magnitude, across the cohort, from 2.2 × 10^−3^ cell^−1^ cell div^−1^ (tumour M) to 1.7 × 10^−2^ cell^−1^ cell div^−1^ (tumour U) (Fig 6A). Median methylation rate spanned 6.5-fold (range: 7.5 × 10^−4^ to 4.8 × 10^−3^ cell div^−1^; Fig 6B), comparable to the fission-rate dispersion, whereas demethylation rates were more consistent (4.1-fold, range: 7.9 × 10^−4^ to 3.3 × 10^−3^ cell div^−1^; Fig 6B, 7). The relatively conserved demethylation rate is consistent with CpG maintenance fidelity acting as a stable cell-biological property, whereas fission rates vary more widely and encode tumour-specific growth histories. Note, however, that the individual posterior 90% credible intervals for fission rate span 1.3–1.7 orders of magnitude within each tumour (Table 3), which is wider than the between-tumour spread of posterior medians; cross-cohort comparisons should therefore be read as ordinal rather than as precise point separations.

**Fig 6.**
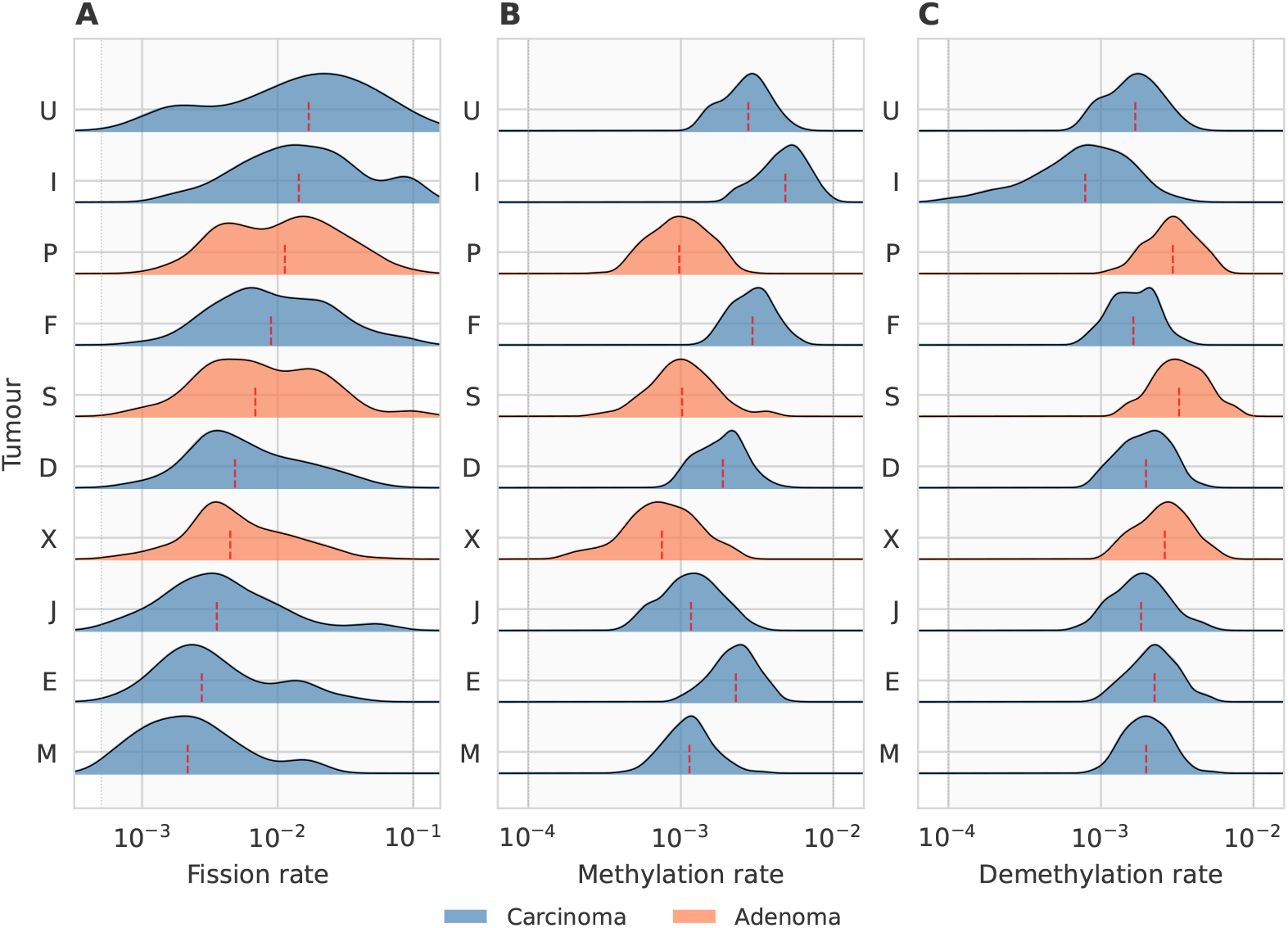
Posterior distributions inferred by *methdemon* for each tumour. **(A)** Fission rate. **(B)** Methylation rate. **(C)** Demethylation rate. Dashed red lines represent the median of the distribution.

**Fig 7.**
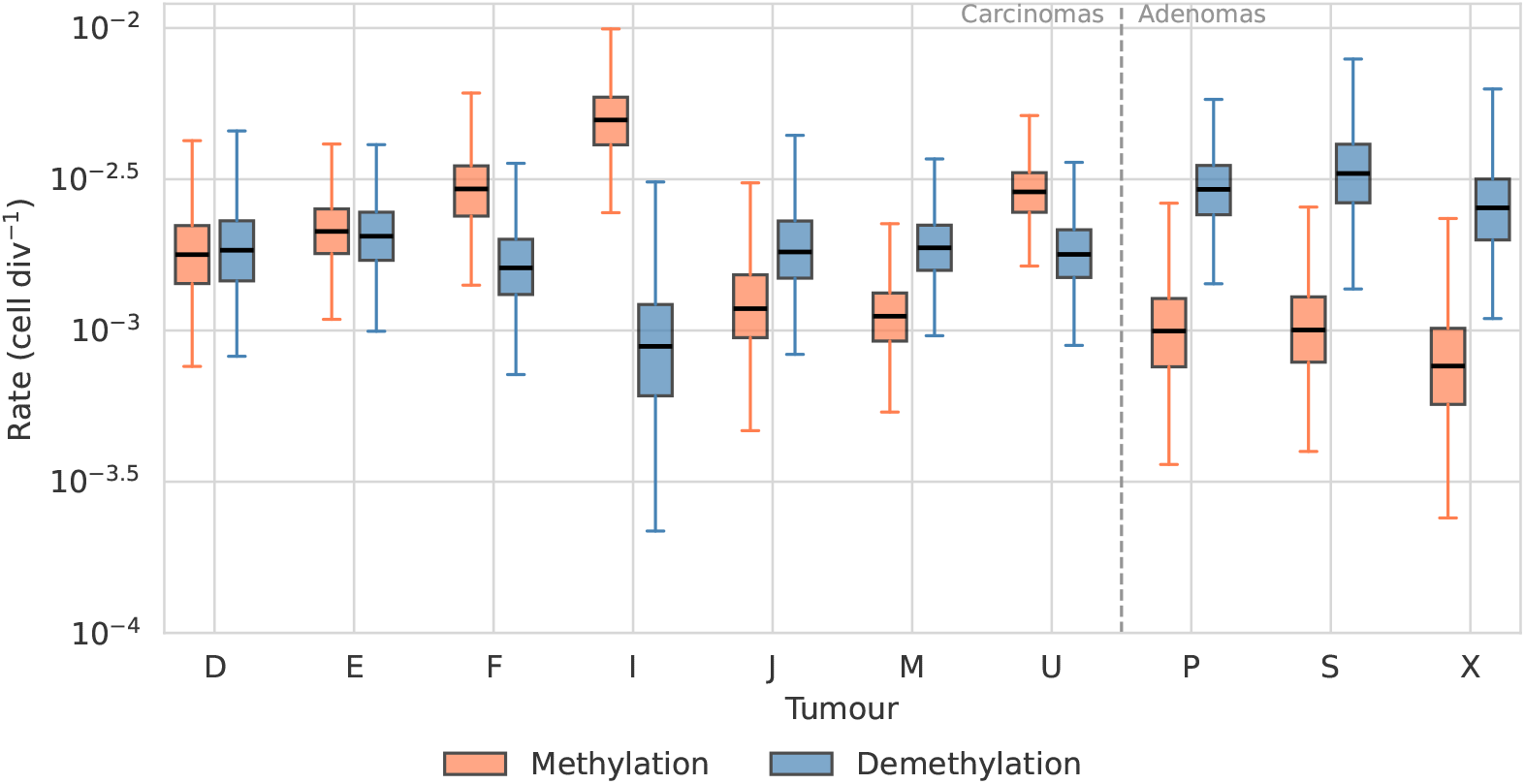
Inferred epimutation rates (methylation and demethylation) across the cohort. Epimutation rates are comparatively conserved across patients (CV = 0.32–0.61).

### Estimation of tumour mitotic ages

Because *methdemon* uses the Gillespie algorithm [25] to schedule events, all inferred rates are in units of (cell division)^−1^, and the framework therefore directly estimates the *mitotic age* of each tumour, i.e. the number of cancer stem cell divisions elapsed during expansion. Multiplying the per-cell fission rate *φ* by the carrying capacity *K* gives the per-gland fission rate Φ = *φK*. Under exponential growth from a single founder gland to *N* glands, the mitotic age is (Methods):

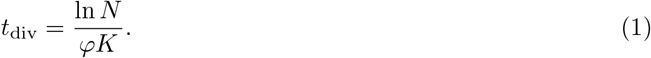

*N* was estimated assuming spherical geometry and a gland cross-sectional area derived from typical colorectal tumour gland dimensions. Full posterior distributions over *t*_div_ were obtained by evaluating Eq 1 at each posterior sample of *φ*.

Median mitotic ages ranged from 10 cell divisions (tumour U) to 73 cell divisions (tumour M) across the cohort (Fig 8). Converting to calendar time requires an estimate of the cancer stem cell division rate, which is not precisely known in colorectal tumours. Using literature estimates spanning once per week to once per month [26, 27], the inferred mitotic ages correspond to tumour expansion times on the order of several months to several years. The wide range across tumours is driven primarily by variation in fission rates, while the uncertainty within each tumour reflects both posterior uncertainty in *φ* and the unknown CSC division rate. Note that the division rate discussed here refers solely to symmetric CSC divisions.

**Fig 8.**
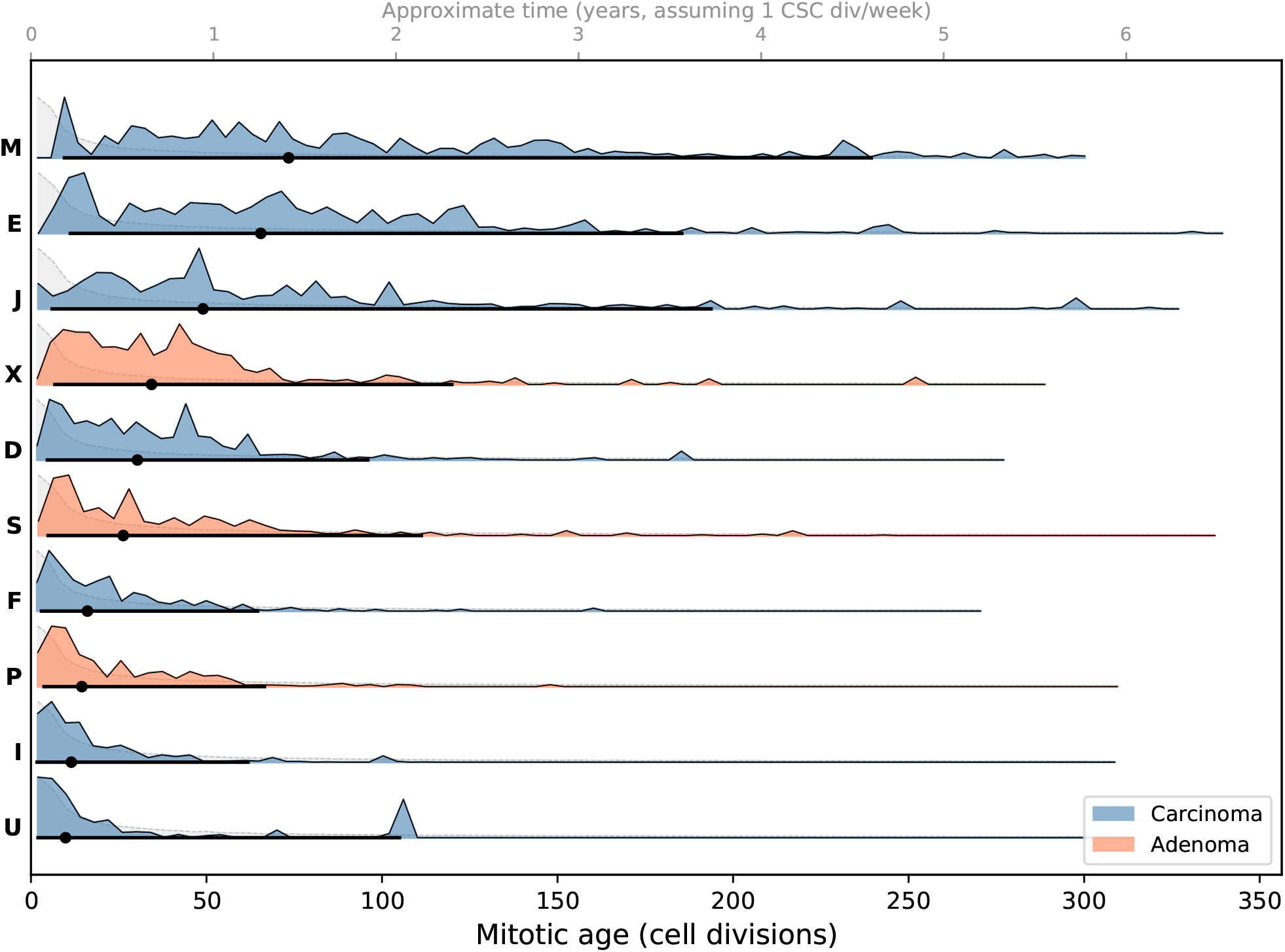
Posterior distributions of tumour mitotic age for each tumour in the cohort. Mitotic ages were computed from the inferred fission rate posterior using Eq 1, with the number of glands *N* estimated from measured tumour diameter. Black dots and horizontal bars indicate posterior medians and 90% credible intervals, and dashed grey curves are the derived prior distributions as per Eq 1.

### Exploratory associations with clinical features

All three adenomas exhibited lower methylation rates and higher demethylation rates than the seven carcinomas (Mann-Whitney *U* = 0, *p* = 0.017 in both cases; Fig S5A), with approximately two-fold differences on average. No significant correlation was observed between inferred fission rate and tumour size (Spearman *ρ* = −0.09, *p* = 0.80), patient age (*ρ* = 0.26, *p* = 0.48), or clinical stage (*ρ* = 0.17, *p* = 0.72, carcinomas only, *n* = 7) (Fig S5B). A trend was observed between patient age and demethylation rate (Spearman *ρ* = −0.56, *p* = 0.093) but this did not reach nominal significance. Neither patient age nor tumour size differed significantly between adenomas and carcinomas in this cohort (Mann-Whitney *p* = 0.68 and *p* = 1.00, respectively), suggesting that confounding by these variables is unlikely, although the sample sizes preclude a definitive assessment. Notably, the wide range of variation in fission rates is preserved when restricting to carcinomas alone (*n* = 7; range 0.0022–0.017; CV = 0.78), showing that this finding does not depend on pooling adenomas and carcinomas. These exploratory associations are based on small, unbalanced groups (*n* = 3 vs *n* = 7) without correction for multiple comparisons, and should be regarded as hypothesis-generating observations requiring validation in larger cohorts.

## Discussion

We have presented a framework for inferring colorectal tumour growth parameters from multi-region bulk fCpG methylation arrays. By combining an agent-based simulation of gland-level tumour expansion (*methdemon*) with ABC-SMC inference (*methabc*), our method for the first time extends fCpG-based inference methods [6–8] to the important use case of solid tumours. Our approach also extends the lineage tracing framework of previous methylation-based colorectal tumour ancestry studies [3–5] in several respects. By explicitly modelling the spatial branching structure of gland fission, we capture the hierarchical patterns that arise from the two-sided sampling design, rather than treating inter-sample distances as exchangeable. Additionally, the use of ABC-SMC provides full posterior distributions over growth parameters, naturally quantifying uncertainty.

Applying our method to multi-region data from 10 colorectal tumours revealed that fission rates vary approximately eight-fold across patients, alongside a comparably dispersed methylation rate (6.5-fold range) and a more conserved demethylation rate (4.1-fold), consistent with previous observations that CpG maintenance fidelity is a relatively stable cell-biological property [1, 2]. Large differences between patients in the extent of intratumour diversity (as measured by fCpG divergence) are therefore driven by variation in both growth dynamics and epimutation rates.

An exploratory association between clinical features and inferred parameters was identified: higher methylation rates and lower demethylation rates in carcinomas (*n* = 7) than in adenomas (*n* = 3). A trend between patient age and demethylation rate was also observed but did not reach nominal significance. Importantly, neither patient age nor tumour size differed significantly between the adenoma and carcinoma groups in this cohort, reducing concern about simple confounding, though multivariate assessment is infeasible for our sample size. Our finding of wide variation in fission rates holds also when the analysis is restricted to carcinomas alone.

No significant correlation was observed between inferred fission rate and tumour size at resection. While this may partly reflect limited statistical power, the result is also consistent with the expectation that tumour size at a single time point is a poor proxy for growth rate: tumours of identical size may have been growing for different durations, and size at resection is confounded by the timing of clinical detection. Moreover, fission rate as inferred here reflects the expansion-phase dynamics, whereas size at resection integrates growth, turnover, and any periods of dormancy or regression.

Several limitations should be noted. Perhaps most importantly, the cohort of ten tumours limits statistical power for detecting associations with clinical features. The study also lacks a baseline control of normal colonic crypts from the same patients. Matched normal crypt methylation profiles from surgical margins would provide a direct patient-specific baseline and confirm that the inferred parameters are specific to tumour growth. Our finding that tumour epimutation rates are considerably higher than those estimated from normal ageing studies [1, 2] is nevertheless consistent with accelerated dynamics during neoplastic expansion and also reflects our selection of fCpG sites that fluctuate over tumour-relevant timescales.

Driver mutation rate and selective advantage proved to be non-identifiable from the available fCpG array data. Hence, while our results are consistent with effectively neutral inter-gland dynamics [14, 15], they do not exclude weak to moderate intra-gland selection. The ability of EVOFLUx to detect subclonal selective sweeps in lymphoid cancers [8] implies potential for denser or targeted within-gland sampling to test for selection in solid tumours.

Our model assumes a well-mixed stem cell population within each gland and does not capture spatial structure within individual glands. While the bulk-sampled arrays make this assumption reasonable, one may wonder about the influence of a gland’s spatial structure on its internal population dynamics. Constraining the stem cell niche to a quasi-one-dimensional population structure at the base of a gland may change the effective population size relative to a well-mixed model of the same census size by influencing the rate of drift and fixation within glands [28]. A more complex and computationally-intensive model would be needed to explore such effects.

Two caveats constrain the interpretation of our inferred mitotic ages. First, our model counts only symmetric stem-cell divisions along the expansion tree. Real colorectal tumour stem cells also divide asymmetrically and cycle at capacity, and because epimutation in the model is coupled to division, these events also generate fCpG switches. The inferred rates *γ*_meth_ and *γ*_demeth_ should therefore be read as effective epimutation rates per symmetric division, analogous to the effective mutation rate per division in neutral models of tumour evolution [15]. Recovering a total-divisions estimate would require knowing the fraction of stem-cell divisions that are symmetric, which is not identifiable from bulk fCpG data. The second caveat is that the converting from cell divisions to calendar time introduces substantial uncertainty because cancer stem cell division rates are not precisely known and may vary across tumours [26, 27]. A rescaling based on estimates of CSC division rates nevertheless places tumour expansion times within a clinically plausible range of several months to several years [14].

The modular design of *methdemon* and *methabc* facilitates adaptation to other cancer types for which spatially resolved methylation arrays can be obtained and appropriate mechanistic models can be constructed. Because the input data are relatively inexpensive, the approach could in principle be applied at cohort scale to ask whether growth history carries prognostic information. The framework could also be integrated with matched genomic data to provide orthogonal validation of inferred growth parameters through comparison with somatic mutation-based phylogenies.

## Methods

### Patient cohort and sample collection

Tumour samples (Table 1) were excess tissues obtained in the course of routine clinical care at the University of Southern California School of Medicine with local Institutional Review Board approval. The tumour samples were examined fresh and approximately 0.5cm^3^ chunks were sampled from opposite tumour sides. From each chunk, single tumour glands (~ 10, 000 cells) were isolated with an EDTA-washout method [29].

### Methylation array processing

Gland DNA methylation (8 glands per tumour) was measured with EPIC bead arrays (Illumina) using the Restore protocol and the manufacturers’ protocols [30]. IDAT files were processed using the noob normalisation function in the minfi R package [31].

### Identification of fluctuating CpG loci

Fluctuating CpG (fCpG) loci were identified as the subset of CpG sites whose methylation state is neither constitutively methylated nor unmethylated across the population, and that show the least coherence across samples, rendering them informative about cell population dynamics rather than cell type identity.

We identified fCpG loci using methylation array data from The Cancer Genome Atlas (TCGA) colorectal cancer cohort, following an adaption of the procedure described in Gabbutt *et al*. [8]. Briefly, we employed the 138 colorectal tumours with purity of at least 0.6 (assessed via ABSOLUTE [32]) and the 38 normal colon tissue samples as our discovery set. We retained CpGs with a high standard deviation across bulk tumour samples, with a mean methylation of approximately 0.5 and with a high Laplacian score (i.e. CpGs that did not preserve the nearest neighbour graph). Unlike Gabbutt et al. [8], we also excluded CpGs that had a high absolute distance from 0.5 in normal colon samples. This procedure identified 1,258 fCpGs specific to colorectal tumours, of which 1,164 were present in all our patient samples and so were used for all subsequent analyses. Using a large reference cohort for locus identification ensures robustness against the sample size limitations of individual patient cohorts.

### *methdemon* : a deme-based model of fluctuating methylation arrays in colorectal cancer

#### Model structure

We developed *methdemon*, an agent-based model written in C++ for simulating the growth of a colorectal tumour and the corresponding dynamics of fCpG methylation arrays across tumour glands. The model represents a tumour as a collection of glands (demes), each containing a population of cancer stem cells with proliferative potential up to a maximum carrying capacity *K*. The well-mixed assumption within each gland is consistent with bulk sequencing of gland samples and reflects the hierarchical organisation of colorectal tumours, in which a small stem cell compartment maintains the gland population [12, 13]. This model structure is inspired by *demon*, a deme-based oncology model [33, 34] used in prior studies of tumour evolution [35–37].

Event scheduling follows the Gillespie algorithm [25], in which a deme is selected first with probability proportional to its total event rate, followed by selection of a cell within that deme. At carrying capacity, cellular dynamics within a gland follow a Moran process, with events consisting of cell birth, cell death, and gland fission. Upon cell division, each methylated fCpG site in the daughter cell undergoes demethylation independently with probability *γ*_demeth_ per site per division, and each unmethylated site undergoes methylation with probability *γ*_meth_ per site per division. Epimutation events are thus coupled to cell division, consistent with the predominantly replication-coupled origin of methylation errors [1, 2].

Driver mutations arise with probability *µ* per cell per division and confer a multiplicative proliferative advantage *s* to the carrier. Each cell tracks its fCpG array as a binary vector of length *L*, and each deme tracks the average fCpG array across its cell population as a vector of methylation fractions.

#### Gland fission

Gland fission is modelled as a neutral spatial branching process, consistent with the predominant mechanism of colorectal tumour growth [10] and with findings supporting neutral inter-gland dynamics [14]. A fission event distributes the gland population randomly into two equal halves, after which each daughter gland repopulates independently to carrying capacity. The per-cell fission probability *ϕ* determines the rate at which new glands are produced.

To focus computational resources on the subset of glands that are ultimately sampled, the model distinguishes between *tracked* and *untracked* fissions. Untracked fissions produce glands that are not followed to the end of the simulation; they contribute to the population dynamics of the tumour but not to the final output. The probability of a fission event being tracked is

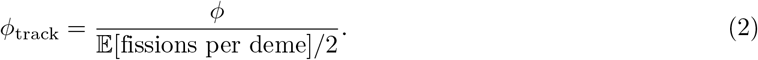

This ensures that the expected number of fission events before a tracked fission is approximately half the mean total number of fissions. Additionally, the distribution of inter-fission intervals at time *t* follows the Poisson distribution with mean 2*ϕt*, as derived in [38].

#### Model parameters and stopping conditions

The full list of model parameters is given in Table 4. The simulation terminates when the mean number of fissions per tracked deme reaches a target value *F*, corresponding to a tumour of *N* = 2^*F*^ glands in expectation. An optional post-growth steady-state turnover phase can be activated, in which cell birth and death continue at carrying capacity without further fission, approximating a growth-saturated regime.

**Table 4.** Parameters of the *methdemon* model.

| Parameter | Description | Units |
| --- | --- | --- |
| $K$ | Deme carrying capacity | cells |
| $\phi$ | Fission rate per cell | $\text{cell}^{-1} (\text{cell division})^{-1}$ |
| $\mu$ | Driver mutation rate | $\text{cell}^{-1} (\text{cell division})^{-1}$ |
| $s$ | Selective advantage of driver mutation | – |
| $\gamma_{\text{meth}}$ | Methylation rate per fCpG site | $(\text{cell division})^{-1} \text{ site}^{-1}$ |
| $\gamma_{\text{demeth}}$ | Demethylation rate per fCpG site | $(\text{cell division})^{-1} \text{ site}^{-1}$ |
| $L$ | Number of fCpG sites per cell | site |
| $F$ | Target mean fissions per deme | – |

### *methabc*: approximate Bayesian computation inference workflow

#### ABC-SMC framework

Parameter inference was performed using approximate Bayesian computation with sequential Monte Carlo (ABC-SMC), implemented on a high-performance cluster with the pyabc package [39, 40]. ABC-SMC iteratively refines a population of parameter particles through successive generations, accepting particles whose simulated outputs fall within a tolerance *ε* of the observed data. The tolerance threshold was determined adaptively at each generation using the `SilkOptimalEpsilon` method [41]. Each generation used 1,000 particles with an acceptance rate of approximately 2%. Inference was run for 9-13 generations until the tolerance threshold and marginal posterior distributions stabilised between successive generations.

All parameters except for selective advantage were drawn from a log_10_-transformed prior to ensure uniform exploration of parameter space across multiple orders of magnitude. Prior distributions are given in Table 2.

The deme carrying capacity was fixed at *K* = 100 for all inference runs, consistent with colorectal tumour stem cell fraction estimates of ~1% [11, 12] given typical gland sizes of ~10^4^ cells. Because the per-gland fission rate enters the model as the product *φK*, the specific value of *K* is a modelling convention rather than a free parameter, and *φK* is the biologically meaningful inferred quantity (§).

#### Summary statistics

Two complementary summary statistics were used to compare simulated and observed fCpG array data.

##### Inter-gland distance matrix

For a tumour consisting of *N* sampled glands with average fCpG arrays **a**_1_, …, **a**_*N*_ ∈ [0, 1]^*L*^, the inter-gland distance matrix **D** is defined as

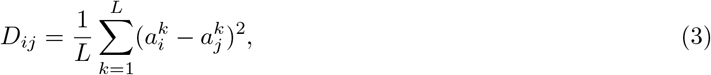

where *L* is the number of fCpG sites. The squared difference emphasises large inter-gland divergences and reduces the influence of small stochastic fluctuations. Two inter-gland distance matrices **D**_1_ and **D**_2_ are compared using the normalised Frobenius norm of their difference:

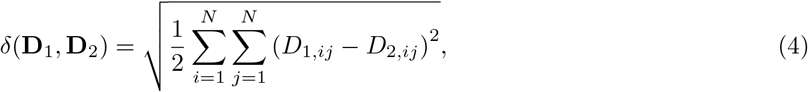

where the factor of 1*/*2 accounts for the symmetry of the distance matrix.

##### Per-gland fCpG distribution distance

To capture information about the shape and bias of individual gland fCpG distributions, which encodes the relative rates of methylation and demethylation, we additionally compute the mean Wasserstein distance between corresponding gland fCpG distributions in the simulated and observed data:

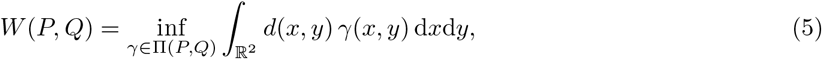

where Π(*P, Q*) is the set of all joint distributions with marginals *P* and *Q*, and *d*(*x, y*) = |*x* − *y* |.

The total distance used for ABC acceptance was a sum of the two statistics:

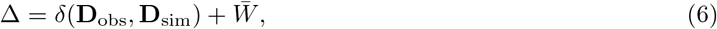

Where 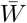 is the mean per-gland Wasserstein distance. The relative contribution of the two components was managed by `pyabc.AdaptiveAggregatedDistance`, which rescales each component at each ABC-SMC generation so that both contribute approximately equally to the acceptance threshold. This adaptive scheme removes the need for a fixed weighting parameter and accommodates changes in scale as the posterior concentrates during successive generations.

#### Posterior predictive checks

We performed posterior predictive checks to assess whether the inferred model adequately reproduces the observed data. For each tumour, 100 parameter sets *θ*^(*i*)^ were sampled from the final-generation ABC posterior distribution. For each sampled *θ*^(*i*)^, one tumour simulation was run using *methdemon* with the same configuration as the corresponding inference run (8 sampled glands, 2 sides, *K* = 100). The simulated inter-gland distance matrix was computed for each replicate, and the distribution of simulated pairwise distances was compared to the observed values. A model fit was considered adequate if the observed mean inter-gland distance fell within the central 90% of the posterior predictive distribution. Pairwise coverage was defined as the fraction of observed inter-gland distances falling within the element-wise 90% posterior predictive interval.

For all 10 tumours in the cohort, the observed inter-gland distance fell within the central 90% of the posterior predictive distribution (Fig S3). Pairwise coverage - the fraction of inter-gland distances falling within the 90% posterior predictive interval - exceeded 96% for all tumours (range: 96–100%). Within-side pairwise distances showed slightly lower coverage (75–100%) than between-side distances (100% for all tumours), suggesting that the model captures the overall magnitude and hierarchical structure of fCpG divergence, though intra-side variation may be underestimated in a minority of tumours.

#### Synthetic parameter recovery

To validate that the inference workflow recovers the identifiable parameters, synthetic parameter recovery experiments were performed. A set of ground truth parameter vectors 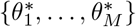 was drawn from the prior distribution, and for each 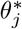 a synthetic tumour was simulated using *methdemon*. The full ABC-SMC inference workflow was then applied to each synthetic dataset to obtain a posterior distribution *p*(*θ* | **D**_*j*_). Recovery was assessed by comparing the posterior median to the ground truth value for each parameter, and by reporting the fraction of experiments in which the ground truth fell within the 90% credible interval (Fig S2). Epimutation rates and the per-gland fission rate *φK* were reliably recovered; *µ* and *s* were not, consistent with the sensitivity analysis described above.

#### Mitotic age estimation

The mitotic age of each tumour was estimated from the inferred fission rate posterior as follows.

Under the Gillespie algorithm, all rates in *methdemon* are expressed per cell division. The per-gland fission rate is Φ = *φK*, where *φ* is the inferred per-cell fission rate and *K* = 100 is the deme carrying capacity. Under exponential growth from a single founder gland, the expected number of glands after *t* cell divisions is E[*N* (*t*)] = *e*^Φ*t*^, giving

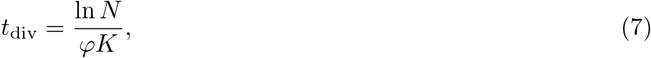

where *N* is the total number of glands in the tumour at resection.

Because gland counts are not directly available, *N* was estimated from the approximate tumour diameter *d* by assuming spherical geometry:

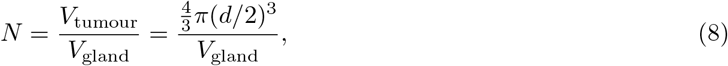

where *V*_gland_ ≈ 1.7 × 10^−3^ mm^3^ was derived from typical colorectal tumour gland dimensions: approximately 70 µm diameter and 450 µm depth [13, 26].

For each posterior sample of *φ* we computed *t*_div_, yielding a full posterior distribution over mitotic age. Conversion to calendar time was performed by dividing *t*_div_ by assumed CSC division rates spanning one division per week to one division per month, following published estimates [26, 27].

The estimator *t*_div_ = ln *N/*(*φK*) should be read as expansion phase symmetric mitotic age: *methdemon* contains only symmetric divisions, so asymmetric and post-expansion contributions are absorbed into the inferred rates rather than resolved as additional divisions.

As an independent cross-check, we also estimated mitotic ages from the within-gland fCpG distributions using a two-state approximation to the Markov chain methylation dynamics described in [2]. For each gland, the mean polarisation, defined as

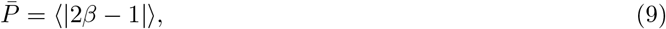

was computed across all fCpG loci (Fig S6A). This quantity measures how far the methylation array has drifted from fully methylated or unmethylated states. Treating each locus as a two-state system in which methylation and demethylation events occur independently at each cell division, polarisation decays geometrically, and the mitotic age can be estimated as

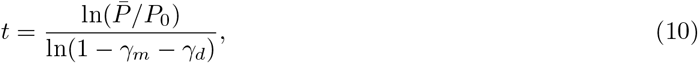

where *P*_0_ is the maximum polarisation observed across the cohort and *γ*_*m*_, *γ*_*d*_ are the posterior median epimutation rates for that tumour (Fig S6B).

#### Turnover sensitivity analysis

To assess whether including steady-state turnover would substantially alter the inter-gland divergence patterns on which inference relies, we performed a grid search across the model’s parameter space at four different lengths of the post-growth turnover phase (0%, 15%, 30%, 50%).

Two hundred parameter sets were drawn from the inference prior via Latin hypercube sampling. For each parameter set, *methdemon* was run at all four turnover fractions using the same random seed, so that the expansion phase was identical and the only difference was the duration of the subsequent turnover period. For each run, the 8 × 8 inter-gland distance matrix was computed and summarised by (i) the mean pairwise distance, (ii) the ratio of between-side to within-side distances, and (iii) the Frobenius distance between the distance matrix at each turnover fraction and the 30% baseline.

#### Statistical analysis

All statistical tests were two-sided with a significance threshold of *α* = 0.05. Given the small cohort size (*n* = 10), effect sizes are reported alongside *p*-values and results are interpreted with appropriate caution regarding statistical power. The within-side versus between-side distance comparison was assessed using a Wilcoxon signed-rank test across the 10 tumours. Spearman rank correlation was used to assess the relationship between inferred posterior median parameter values and continuous clinical variables (tumour size, patient age). Mann-Whitney *U* tests were used to compare inferred parameter values between categorical groups (sex, adenoma versus carcinoma). Consistency of inferred epimutation rates across patients was quantified using the coefficient of variation (CV) of the posterior medians. All statistical analyses were performed in Python 3.12 using `scipy v1.17.1 (scipy.stats` module).

## Data availability

All fCpG arrays used in the paper are publicly available at https://github.com/vesmanojlovic/crcfcpg.

## Code availability

The agent-based model *methdemon* is available at https://github.com/vesmanojlovic/methdemon. The ABC-SMC inference wrapper *methabc* is available at https://github.com/vesmanojlovic/methabc.

Analysis scripts for posterior extraction, posterior predictive checks, sensitivity analyses, and figure generation are available at https://github.com/vesmanojlovic/crcfcpg.

## Competing interests

C.G. is named on a method to measure evolutionary dynamics in cancers using DNA methylation (GB2317139.0). V.M., D.S., and R.N. declare no competing interests.

## Acknowledgements

We are grateful for the use of City St George’s Hyperion cluster to run the ABC inference workflows for this study. V.M., D.S., and R.N. were supported by the National Cancer Institute of the National Institutes of Health under Award Number U54CA217376. The content is solely the responsibility of the authors and does not necessarily represent the official views of the National Institutes of Health. C.G. acknowledges support from Cancer Research UK (EDDPMA-May23/100059) and the Thornton Foundation. Support was provided to C.G. by Schmidt Sciences, LLC.

## Supporting information

**Fig S1.**
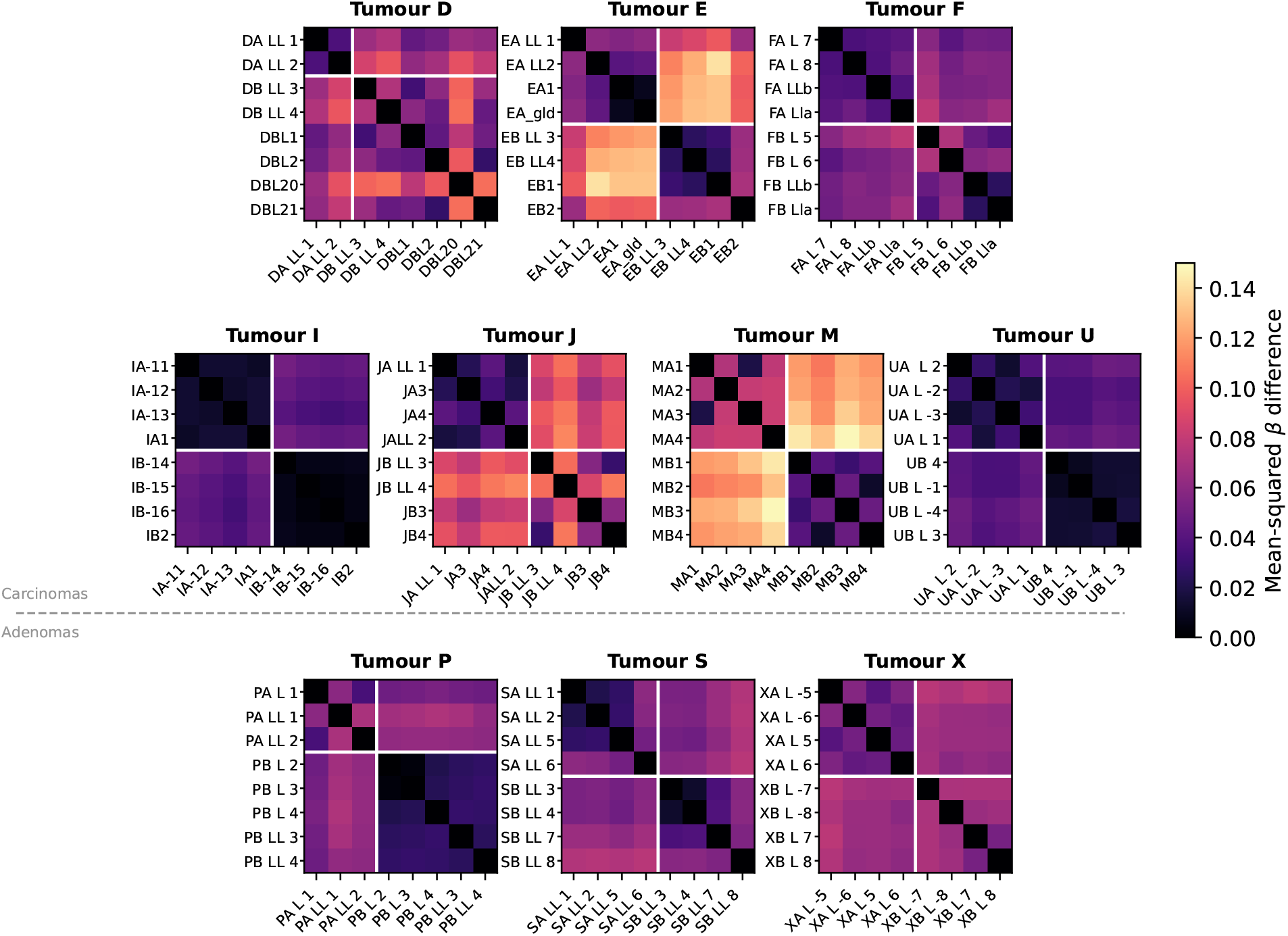
Inter-gland fCpG distance matrices (mean squared difference) for all 10 tumours. Bold lines separate glands from side A (top-left block) and side B (bottom-right block). Block structure is visible in all tumours, with strongest separation in tumours I, U, and M.

**Fig S2.**
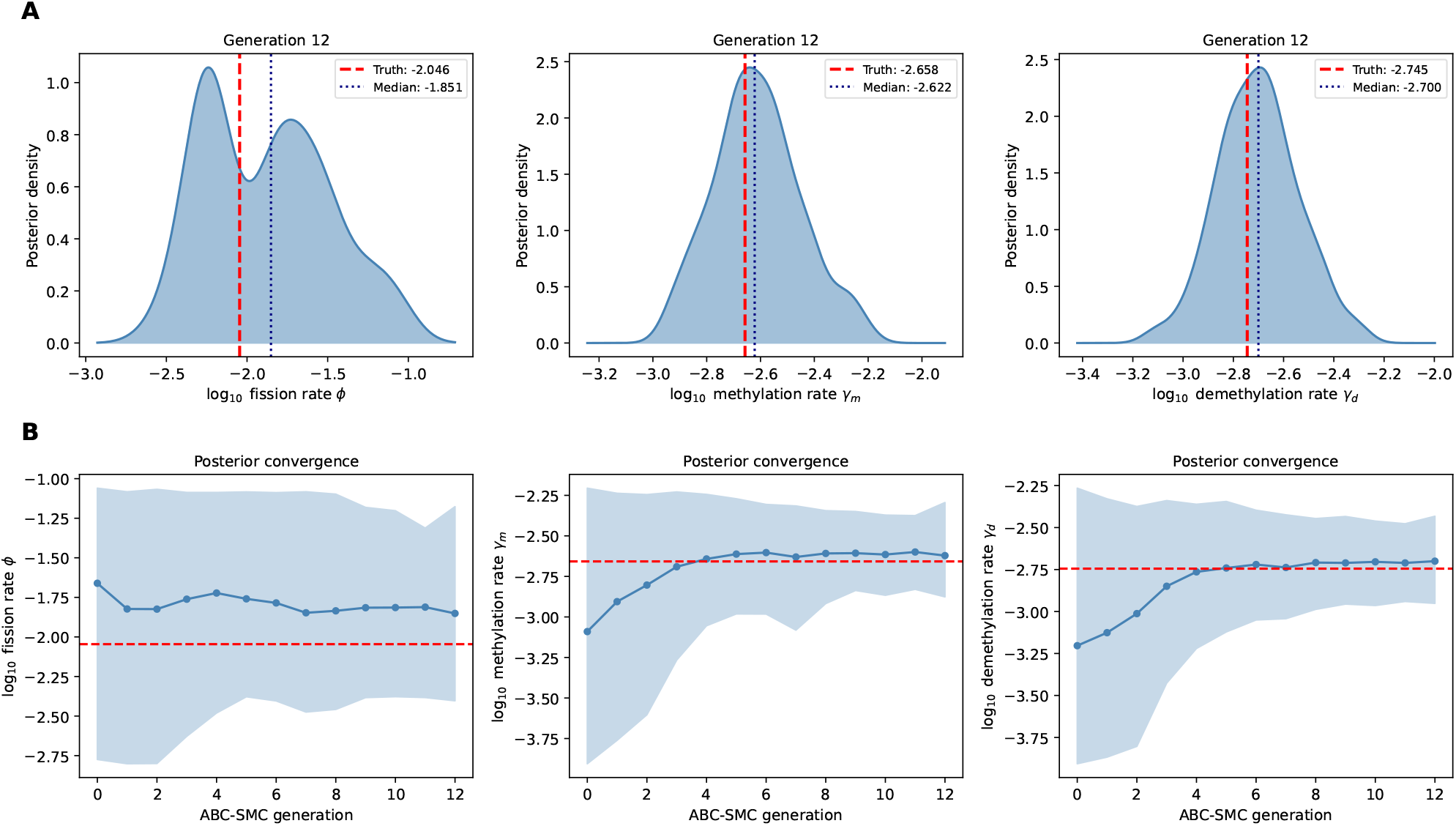
Synthetic parameter recovery. **A:** Final-generation posterior distributions (blue) for the three identifiable parameters, with ground truth values (red dashed lines). All three parameters are recovered accurately. **B:** Convergence of posterior medians and 90% credible intervals across ABC-SMC generations.

**Fig S3.**
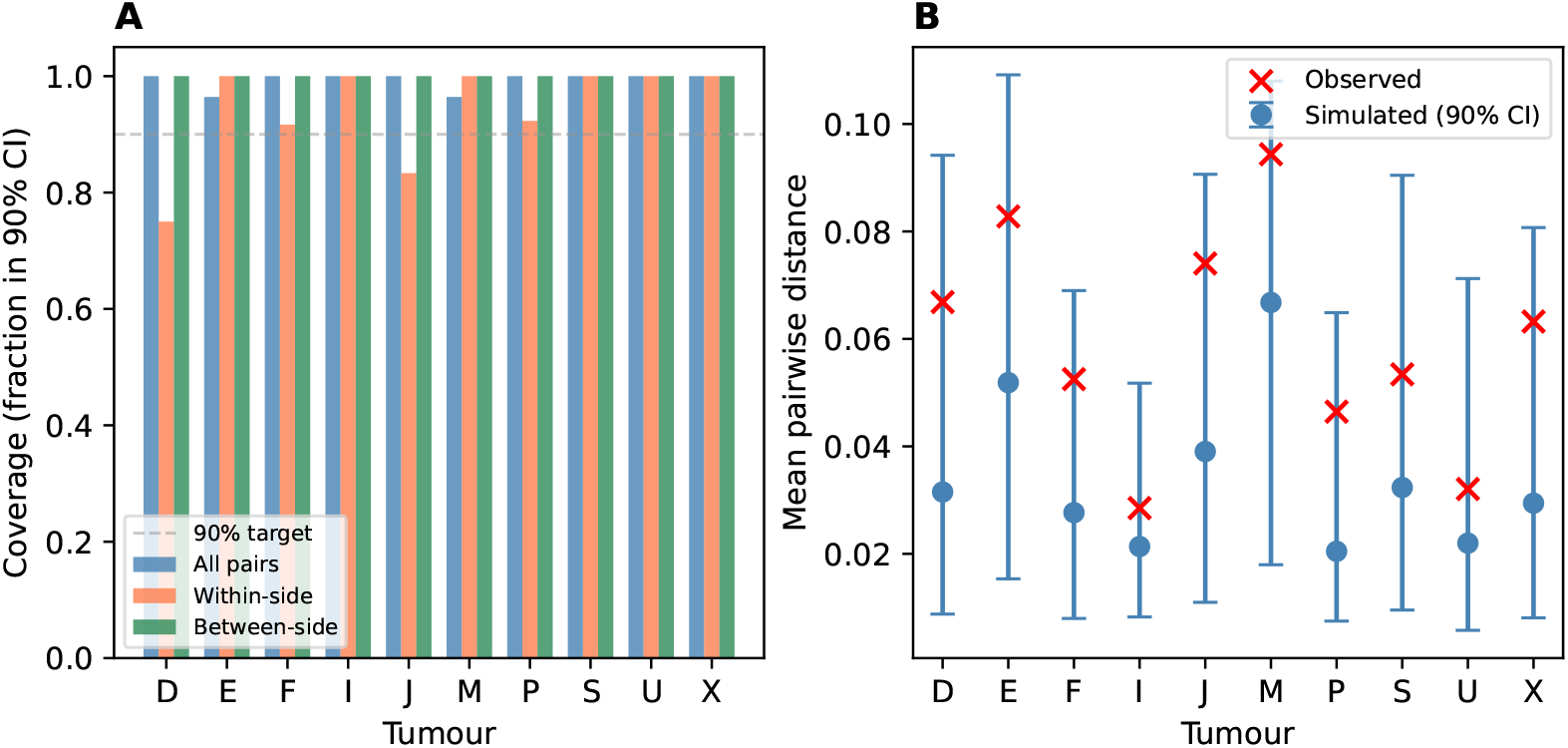
Posterior predictive check summary across the cohort. For each tumour, 100 datasets were simulated from the inferred posterior. **A:** Pairwise-coverage plot of the posterior-drawn simulations. **B:** Observed mean inter-gland distances. Observed values fell within the central 90% of the posterior predictive distribution for all 10 tumours.

**Fig S4.**
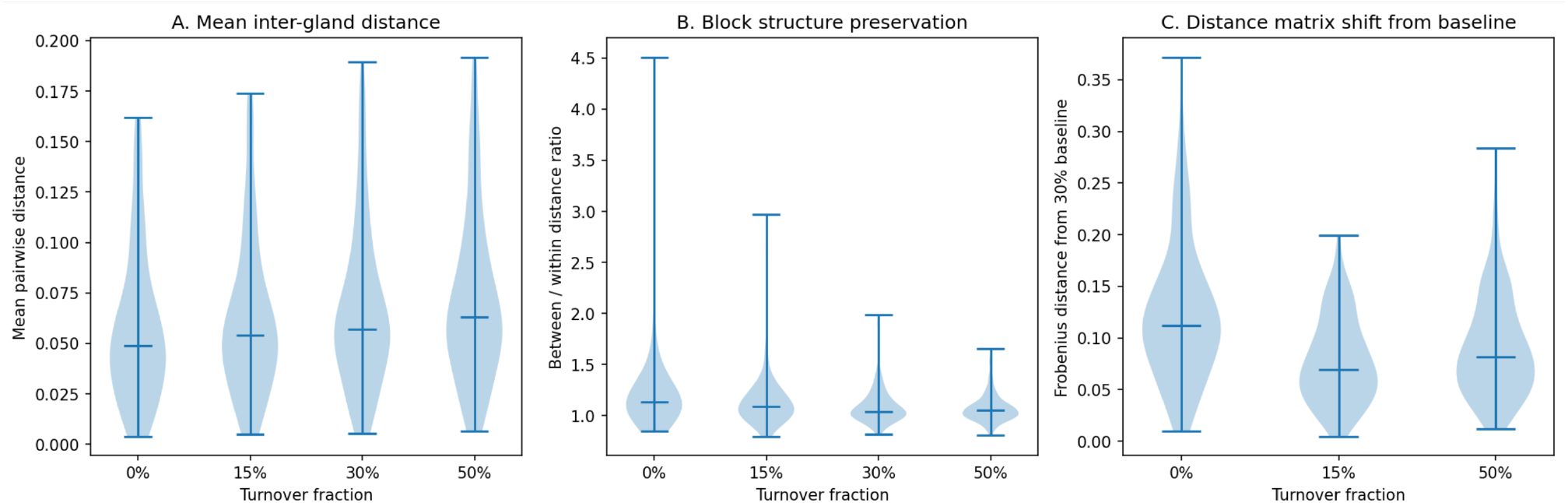
Model sensitivity of *methdemon* to the time spent in turnover after the growth phase as a fraction of the time spent in the growth phase. **A:** mean pairwise inter-gland distance across different levels of post-growth turnover. **B:** The ratio of the inter-gland distances for near and distant glands across different levels of post-growth turnover. **C:** The shift of the distance matrix across levels of post-growth turnover as compared to a 30% turnover fraction.

**Fig S5.**
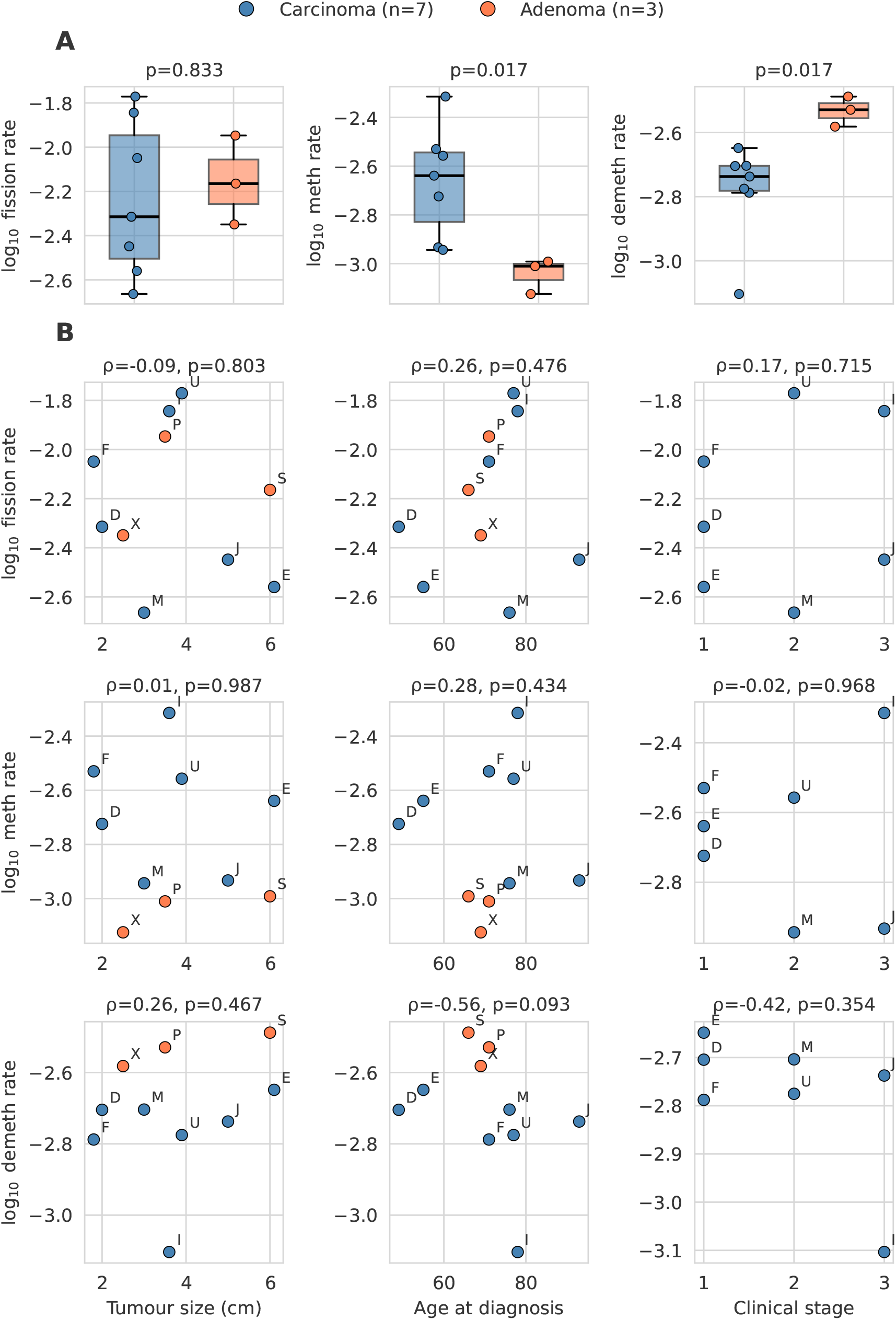
Analysis of how inferred parameter values vary between tumours. **A:** Fission, methylation, and demethylation rate comparison between carcinomas and adenomas in the cohort. **B:** Spearman correlations between inferred parameter values and clinical features.

**Fig S6.**
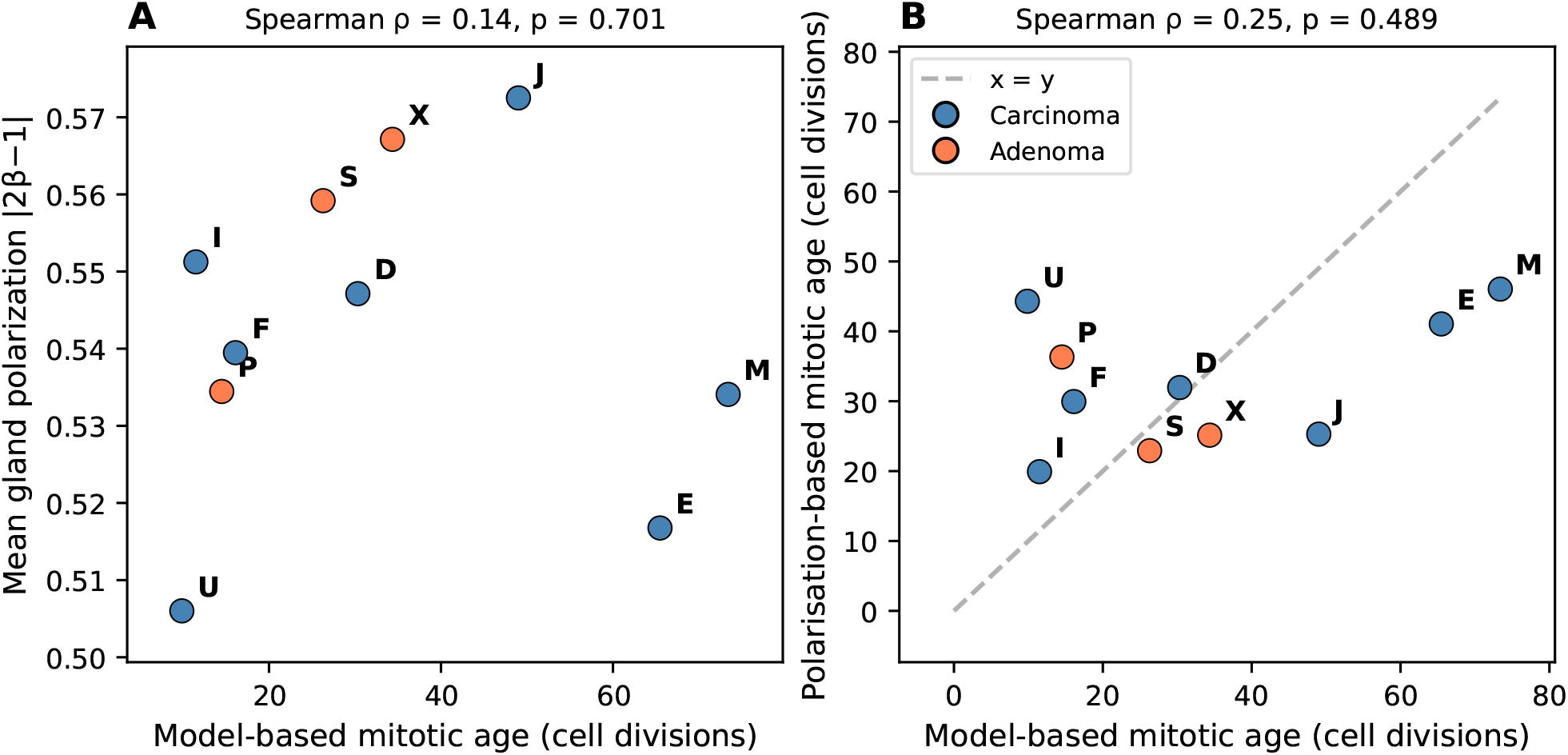
Consistency of mitotic age estimates. **A:** Mean gland polarisation versus model-based mitotic age. **B:** Polarisation-based versus model-based mitotic age estimates.

## Notes

https://github.com/vesmanojlovic/crcfcpg

### Summary of Updates

Posterior predictive checks now discussed only once instead of three times. Figures have been reordered and some figures moved to supplementary. Figure 4 (previously Figure 2) has been simplified. Added Author Summary and Competing Interests. Removed a couple of duplicated references. General polishing of the text.

